# Microscopy Nodes: versatile 3D microscopy visualization with Blender

**DOI:** 10.1101/2025.01.09.632153

**Authors:** Oane Gros, Chandni Bhickta, Granita Lokaj, Yannick Schwab, Simone Köhler, Niccolò Banterle

## Abstract

Effective visualization of 3D microscopy data is essential for communicating biological results. While scientific 3D rendering software is specifically designed for this purpose, it often lacks the flexibility found in non-scientific software like Blender, which is a free and open-source 3D graphics platform. However, loading microscopy data in Blender is not trivial. To bridge this gap, we introduce Microscopy Nodes, an extension for Blender that enables the seamless integration of large microscopy data. Microscopy Nodes provides efficient loading and visualization of up to 5D microscopy data from Tif and OME-Zarr files. Microscopy Nodes supports various visualization modes including volumetric, isosurface, and label-mask representations, and offers additional tools for slicing, annotation, and dynamic adjustments. By leveraging Blender’s advanced rendering capabilities, users can create high-quality visualizations that accommodate both light and electron microscopy. Microscopy Nodes makes powerful, clear data visualization available to all researchers, regardless of their computational experience, and is available through the Blender extensions platform with comprehensive tutorials.

## Introduction

Biology is three-dimensional and dynamic but most representations are limited to two dimensions. The informative and correct mapping from 3D space to 2D visualization, ‘rendering’, is therefore essential for communicating biological 3D data. Multiple scientific software solutions exist for rendering data in a 3D environment, both proprietary, such as Imaris^1^, Arivis^2^ and Amira^3^, and open-source alternatives such as BigVolumeViewer^4,5^, Agave^6^ and Napari^7^. However, these scientific tools are often limited to workflows for specific imaging modalities where editing the rendering pipeline or 3D scene for custom pipelines remains challenging. By contrast, non-scientific 3D rendering software has had a large user base and need for diverse features for a long time, leading to comprehensive and mature software packages in both the proprietary and open-source space.

One of these non-scientific solutions for 3D rendering is the powerful software Blender^8^ which is committed to staying free and open-source long-term. As a generic graphics toolkit, Blender does not specifically support easy loading of microscopy data, nor can it readily handle the large data size of microscopy volumes. Although Blender has been used previously by microscopists, it was only accessible to users with extensive knowledge of both Python and the Blender user interface (for example, Hennies et al.^9^). Here, we present Microscopy Nodes, a bridge between the microscopist and advanced data representation in Blender. Microscopy Nodes seamlessly integrates microscopy data into an ecosystem providing powerful tools for beautiful presentation-quality visualizations, enabling more effective communication of 3D biological data.

## Results

Using Microscopy Nodes, we can load a large variety of microscopy modalities, and target a diverse set of output visualizations. Here, we showcase a selection of different target visualizations that can be achieved with Microscopy Nodes and Blender. The input modalities tested include real-time imaging (Fig 1A, https://uk1s3.embassy.ebi.ac.uk/idr/zarr/v0.4/idr0052A/5514375.zarr), larger datasets, such as a 49 GB expansion microscopy stack (Fig 1B, https://s3.embl.de/microscopynodes/RPE1_4x.zarr, generated for this showcase) and a 14.5 GB electron microscopy data (Fig 1C, EMPIAR-11399). These datasets are hosted publicly in archives such as the IDR^10^, Bioimage archive^11^, and EMPIAR^12^. When available, OME-Zarr datasets from these repositories can be loaded from their address (URI) in Microscopy Nodes. This feature enables interested users to make use of Microscopy Nodes even without their own dataset.

**Fig 1.**
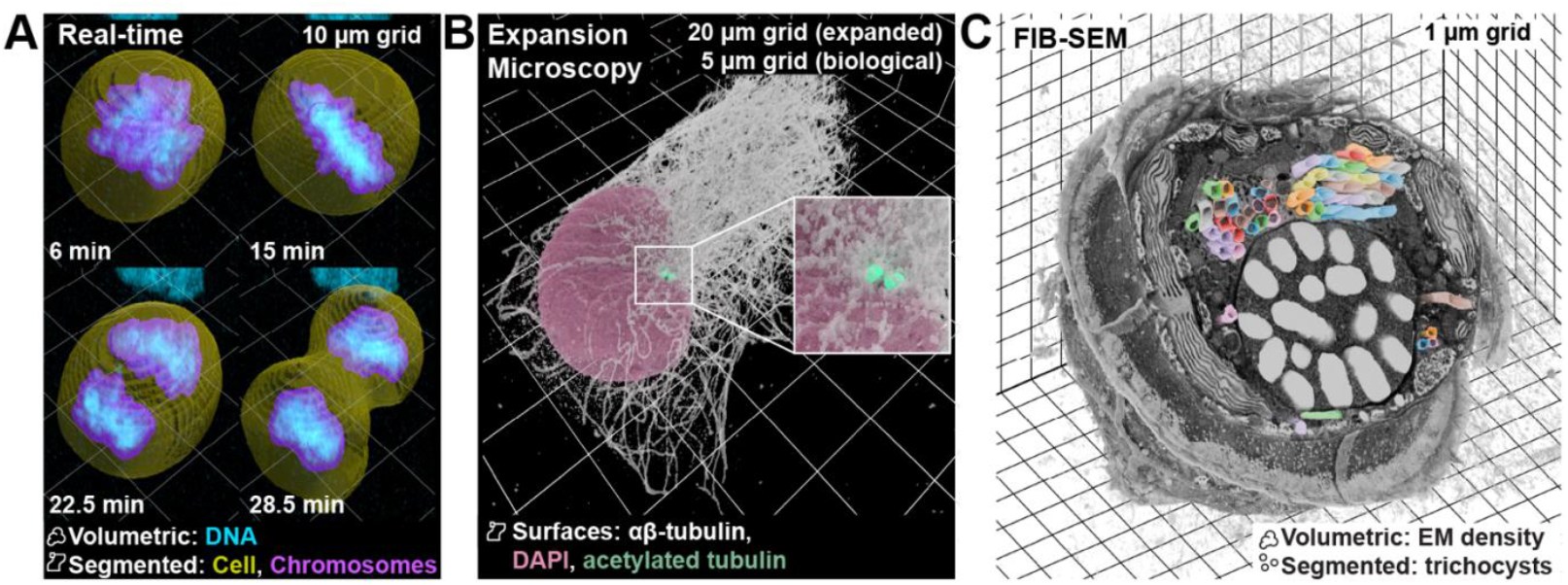
Microscopy Nodes supports diverse data modalities and output formats. **A)** *A time-series of a mitotic cell with segmentations*. The render shows the dynamics of chromosomes (DNA) during mitosis by combining mesh and volumetric rendering modes of data from Walther et al.^25^ **B)** *An isosurface render of a RPE1 cell shows its intricate cytoskeleton*. The expansion microscopy image of the cell is shown as isosurfaces with the αβ-tubulin slightly transparent to highlight the centrioles (acetylated tubulin). The versatility to independently adapt separate channels allows for better communication of the data. **C)** *Blender allows for contextual EM visualization*. A dinoflagellate cell imaged with FIB-SEM and its segmented trichocysts^13^. With Microscopy Nodes and Blender, we can reliably visualize the ultrastructure of cells and show EM data where electron-sparse regions are transparent, with segmentations in the same view. Here the segmentations are cropped to a separate region than the EM density, to show them protruding from the volume.

When loading data with Microscopy Nodes, multiple representations are available, and which representation is appropriate depends on the type of data. For example, the real-time imaging dataset shown here (Fig 1A, Supplemental Video SV1) consists of fluorescent channels and segmentations of one cell and its chromosomes. Here, we visualize one of the fluorescent channels (DNA) as a volumetric and emissive render, meaning that each voxel emits light with an intensity relative to its measured value in the dataset. This way of handling fluorescent data matches the acquisition technique by representing the light-emissive fluorescent probe as a light-emissive signal in the 3D render. In the same view, we included the segmentations as semitransparent surface renders. These options allow us to combine the data and its analyzed interpretation in the same representation with Microscopy Nodes.

The expansion microscopy dataset (Fig 1B) used here has multiple fluorescent channels representing microtubules, acetylated tubulin (used as a centriole marker), and the nucleus. This dataset is challenging to visualize as the microtubule network is very dense, and the difference in scale between the centriole and the cell is large. To emphasize the intricate network of microtubules, we used an isosurface render, where a surface mesh is generated at a single threshold value in the image. The threshold for the surface mesh can be applied on load with Microscopy Nodes and interactively changed in Blender. By using the isosurface, the fibers in the microtubule signal are separated and the differences in height are enhanced compared to a direct volumetric render. Similarly, the centrioles and the nucleus are also shown as isosurface renders. However, to emphasize the small centriole in this larger dataset, we used the interactively editable render properties to add a light-emission property to their surface. Another way to emphasize small structures is to use Blender’s extensive camera controls and animate the visualization properties of the different channels through animation time (Supplemental Video SV2). In this video, we also highlight another key feature of Blender powered by Microscopy Nodes: the ability to combine microscopy data with any 3D illustration in the same 3D environment. Blender has many programmatic and manual options to create 3D illustrations, which we used here to incorporate a model of the centriole and its components into the data visualization (Supplemental Video SV2).

With Microscopy Nodes, we are able to load both light and electron microscopy data, changing only one load setting. To illustrate this, we use a manually segmented dinoflagellate phytoplankton volume EM stack, gathered with Focused Ion Beam Scanning Electron Microscopy (FIB-SEM)^13^. Here, we rendered the data as a volumetric density, which is light-scattering rather than light-emissive. This light-scattering render mode reflects the data acquisition of scanning electron microscopy, as this method captures the scattered electrons of the block surface. Additionally, we load the segmentation of the trichocysts (thread-like organelles that can be ejected) with Microscopy Nodes, which separates the segmentations for each organelle into separate surfaces and applies a colormap per object identity. The color can also be edited per object identity. Additionally, to show the trichocyst segmentation more clearly in the dense EM signal, we use two different slicing regions for the segmentation and the data, showing the trichocysts protruding from the EM density.

### Microscopy Nodes as a tool for effective 3D visualization

With the adaptability of the volumetric rendering of Blender powered by Microscopy Nodes, we are able to communicate new biological results more effectively. To demonstrate this with a novel dataset, we present a late mitotic, dinoflagellate phytoplankton imaged with volume electron microscopy (FIB-SEM). This sample was collected in the environment and cryopreserved by high-pressure freezing. Mitosis in most dinoflagellates is unique, and termed “dinomitosis”^14,15^. It is a form of closed mitosis, where the nuclear envelope remains intact and instead invaginates to form “nuclear tunnels”. These tunnels allow the mitotic spindle to traverse the nucleus and assist with chromosomal segregation without entering the nucleoplasm^16^.

FIB-SEM data is typically presented as single 2D slices of the data (Fig 2A), where in this case, the nuclear tunnels appear as invaginations into the nuclear envelope within the slice plane. This visualization is also supported in Blender, allowing for arbitrary selections of viewing angles (Fig 2B). However, we can also utilize the advanced rendering engine to show these nuclear tunnels in 3D (Fig 2C). To this end, we only show electron densities filtered to the density corresponding to membranes, and rendered all other EM density as transparent (analogous to Fig S1). This approach allows us to look through the dense nucleus and visualize the 3D network of these nuclear tunnels in a single visualization. Utilizing the same method, we can also add a second color for EM densities corresponding to the chromosomes, and show both structures of interest in the same visualization without manual segmentation (Fig 2D), although here the chromosomes obscure some tunnels in the nucleus.

**Fig 2.**
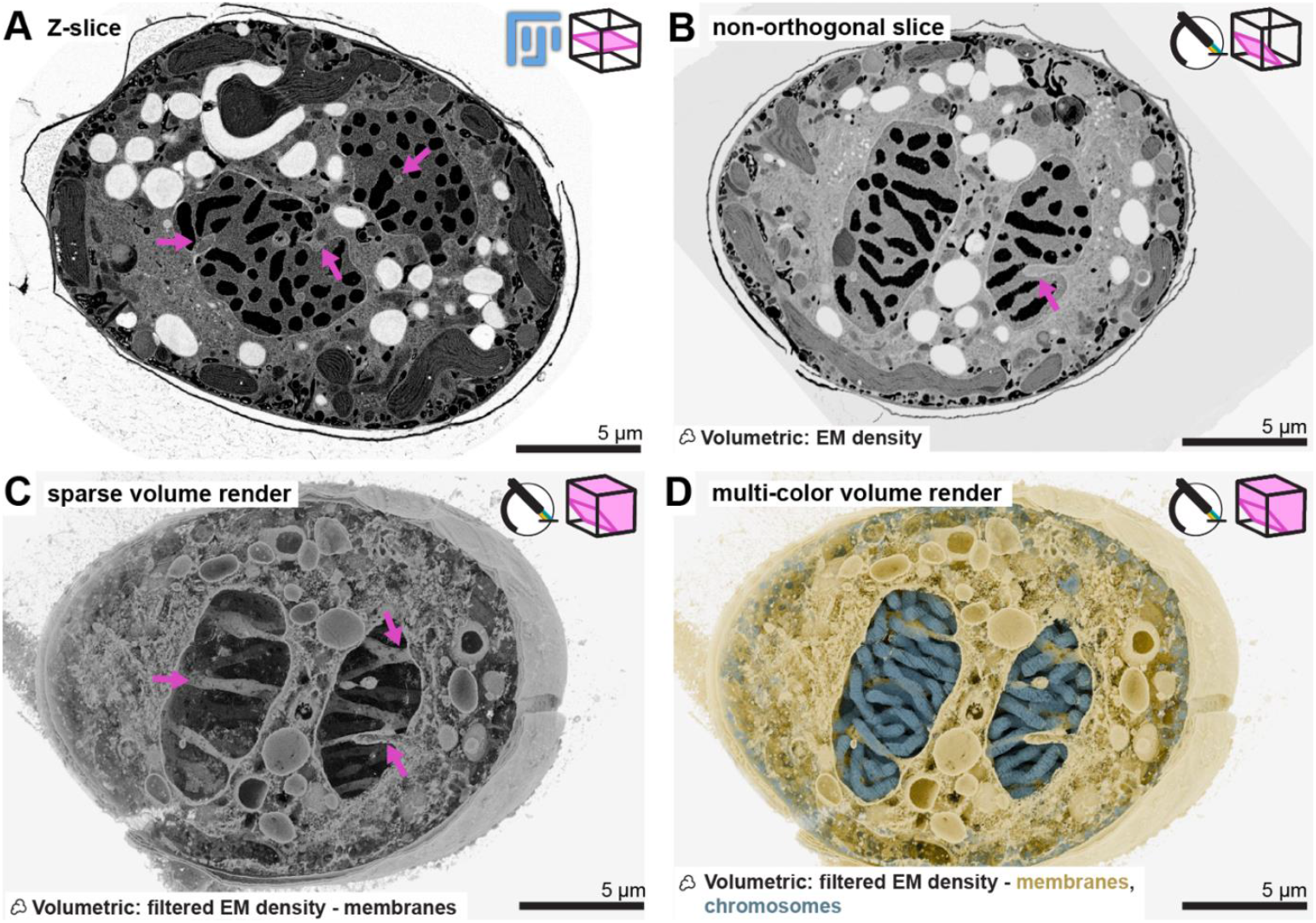
Sparse 3D rendering assists in communicating new biological information. Different renders of the same dinoflagellate cell. Logos in the top right indicate the software used to render (Fiji or Microscopy Nodes), and a cartoon illustrating which data is used in the render (not to scale). Scale bars are shown as bars instead of grids as all images are orthographic projections. **A)** *Z-slice of a late mitotic dinoflagellate showing nuclear tunneling*. A single slice of a FIB-SEM stack of a dinoflagellate showing invaginations in the nuclear envelope (magenta arrows), black is electron dense matter. **B)** *A dense render of an arbitrary slicing axis in Blender of the same dataset*. Here, the data was loaded into Blender with Microscopy Nodes to select a non-orthogonal slicing axis to show the nuclear tunnels (magenta arrows). **C)** *Rendering some EM densities as transparent can give a 3D overview of biological structures*. A volumetric render in Blender, highlighting low EM densities as dense volumetric renders to highlight low-density structures such as membranes. This visualization shows the 3D network of tunnels in the nucleus (magenta arrows) in a single rendered image. **D)** *Splitting the color map highlights different structures*. A 3D render, where EM densities corresponding to membranes are colored yellow, and EM densities corresponding to chromosomes are colored blue. This render shows how we can use selective rendering of certain densities to highlight different structures in dense 3D data. However, this visualization occludes the nuclear tunneling phenotype partially.

As shown here, the best 3D representation of a dataset is dependent on the data that is visualized. Microscopy Nodes offers a single toolkit that allows for very versatile handling of a wide range of data types. Since it is built in the popular and extremely capable toolkit Blender, it also provides an extendable environment for customizing complex target visualizations.

## Design and Implementation

Microscopy Nodes is a Blender extension written in Python, and provides an interface to easily load any microscopy data, giving choices between preset load settings. The user can then interactively use the extensive Blender interface to edit, annotate, animate their data, and render the final output image either from the interface or the command line (Fig 3A). Microscopy Nodes loads microscopy data, saves the data into Blender-compatible files (namely VDB^17^ volume files and alembic^18^ mesh files), loads these files into Blender, and applies presets and Microscopy Nodes-specific user settings. The re-saving into local files means that a local copy of the rendered data must be present on disk, although only the data shown in a single timepoint needs to fit comfortably (2-3x original size, minimally 16 GB) in RAM.

**Fig 3.**
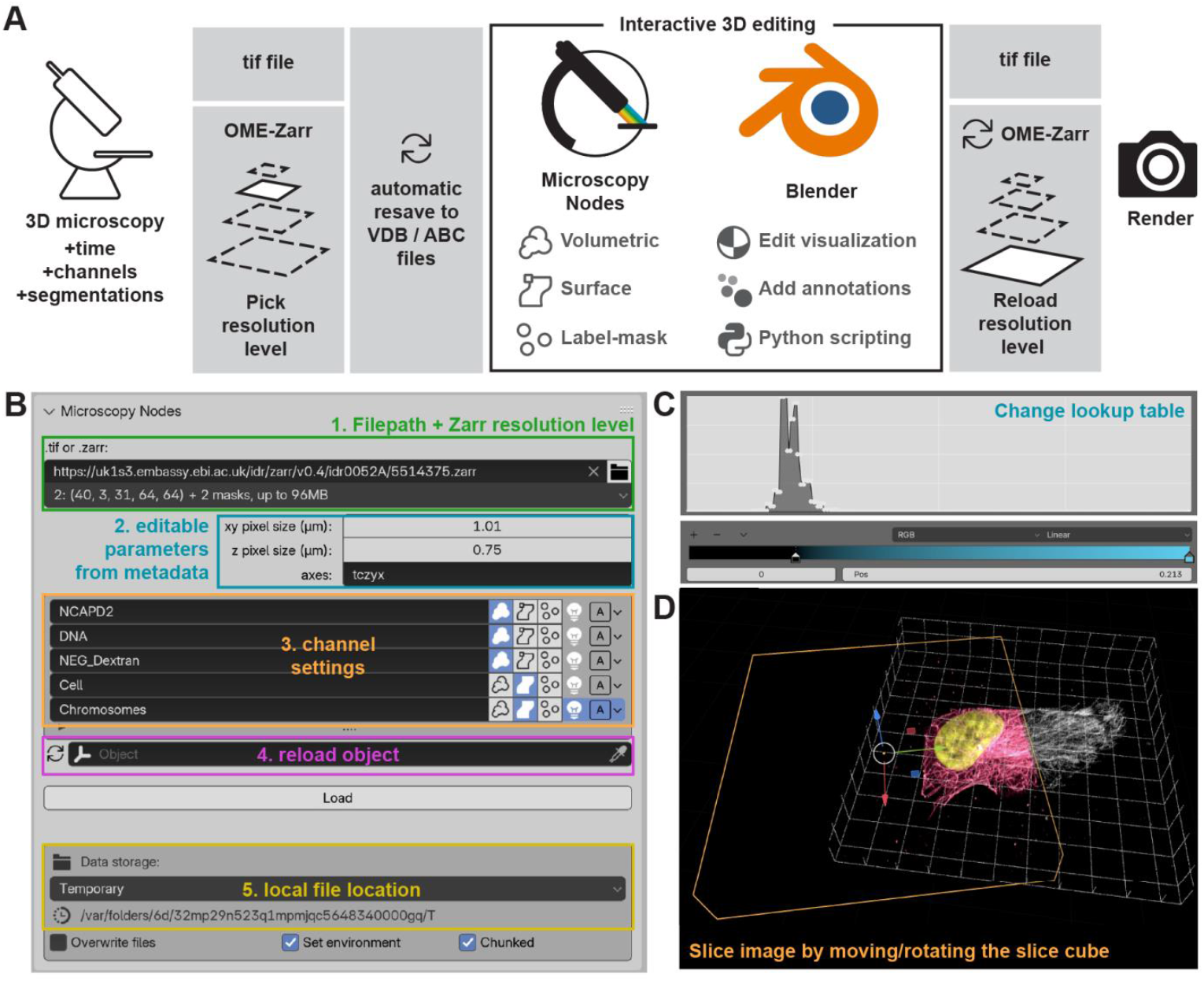
User interface of Microscopy Nodes. **A)** *The Microscopy Nodes workflow allows for interactive use of big data*. Up to 5D microscopy files with segmentation channels can be loaded into the Microscopy Nodes add-on, which resaves the data into Blender-appropriate formats. The Microscopy Nodes Blender interface allows for adjusting settings for each channel and interactive editing. When using pyramidal Zarr data, a smaller dataset can be used interactively, to then up-scale for the final render. Icons used are from the Blender icon library. **B)** *The Microscopy Nodes panel in Blender allows specifying a large variety of image features quickly*. (1.) Input is specified as path to a tif or OME-Zarr file, or using an url linking to an OME-Zarr file in a repository. (2.) Pixel sizes, axis order, and channel names are read out automatically, but they can also be specified manually. (3.) For each channel, users can select (from left to right): load as volumetric, load as isosurface, load as integer label mask, whether the data should emit or absorb light, and surface resolution, respectively. (4.) If specified, Microscopy Nodes will also read out channel names from the metadata. Additionally, users can reload a previous volume to change the underlying data or data resolution, and set environment variables. (5.) By default, Microscopy Nodes will store local files in a temporary folder, but this location can also be specified by the user. All buttons have hover-tooltips in the software. **C)** *Lookup tables can be interactively remapped with a histogram of the voxel values*. Up to 32 colors can be specified in between, with default loading as a linear LUT from black to one color per channel. **D)** *Slicing a volume can be achieved by moving the slice cube in the 3D space*. To illustrate slicing, two copies of the same dataset were loaded, where one is sliced with the selected slice cube (orange, sliced data in magenta), and the other shown unsliced in white.

Microscopy Nodes supports both Tif and OME-Zarr^19^ files, and it can use the pyramidal Zarr data structure to load and reload data from a Zarr URI (optionally in an S3 bucket) at different resolutions. For large OME-Zarr files, a user can load a smaller version of the data to tune the visualization settings interactively. The data can then be reloaded with a higher resolution only for the final render, optionally from the command line. This reloading architecture allows users to handle much larger data than their personal computer can handle interactively, and a project can thus be transferred to a machine with more RAM.

With loading, the user can select specific settings for visualizing the data: from basic options such as showing it as fluorescence (emitting light) or electron (scattering light) microscopy (Fig 3B), color and intensity adjustments (lookup table mapping) (Fig 3C), and slicing (Fig 3D), down to granular control of how light in the scene interacts with the volumetric data (Supplemental Note 1). Microscopy Nodes provides an extensible and versatile environment for 3D microscopy visualization, while also providing useful presets and easy and intuitive tools. By default, the Microscopy Nodes toolkit only uses a small segment of the extensive Blender user interface. However, the vast rest of the interface remains accessible and can be used as needed. As such, Blender allows for, e.g. the drawing of annotations manually through the many modeling tools or programmatic creation and animation of meshes, either from its embedded graphical programming language or Python scripting. Blender can also be further extended with other (scientific) add-ons such as Molecular Nodes^20^ or Mastodon cell-tracking^21^.

## Discussion

With the great versatility in both input modality and target visualization, we believe Microscopy Nodes has the potential to revolutionize the visualization of 3D microscopy data, as it makes Blender, a comprehensive rendering and modeling tool, accessible for all labs and any biologist regardless of their computational experience. This paves the way for Blender to act as a platform for an ecosystem of tools for microscopy data visualization and analysis.

Currently, Microscopy Nodes purely serves a visualization purpose, and the core of the add-on is loading data that is easily adaptable rather than providing analysis tools. However, Microscopy Nodes can already be used as a loading interface for mesh analysis in Blender^22^, and once volumetric math and mesh analysis in Blender mature, we anticipate that new image analysis tools will be incorporated or built on top of Microscopy Nodes.

Microscopy Nodes is available for download directly from the Blender interface, the Blender extensions platform, and the GitHub. It is documented on https://oanegros.github.io/MicroscopyNodes/ and through a series of YouTube tutorials, that are kept up to date with Blender and Microscopy Nodes updates.

## Code availability

Microscopy Nodes is open source and published under a GPL-3 license, hosted at https://github.com/oanegros/MicroscopyNodes, and directly installable from the Blender extension panel.

## Methods

### Electron microscopy sample preparation

In the context of the TREC expedition (https://www.embl.org/about/info/trec/), natural phytoplankton communities were sampled with a 10 µm plankton net near the Kristineberg Centre for Marine Research and Innovation and fractionated with a 40 µm sieve. Making use of the Advanced Mobile Laboratory (AML) deployed at the marine station, the sample was concentrated on a 1.2 µm mixed cellulose ester membrane, and allowed to sediment before high-pressure freezing (EM-ICE, Leica Microsystems). A 1,2 µl drop of the concentrated mix was loaded into a type A carrier (gold-coated copper), 3mm wide and 200 µm deep), and sandwiched with a type B aluminium carrier. Sample was freeze substituted (EM-AFS2, Leica Microsystems), following protocols previously described in literature ^13,23^ . The sample was incubated in 0.1% uranyl acetate dissolved in dry acetone at -90°C for 72 hours. The temperature was then gradually increased to -45°C at 2°C per hour, followed by incubation at -45°C for 10 hours. Subsequently, the sample was infiltrated with increasing concentrations of Lowicryl HM20 resin (10%, 25%, 50%, 75% and 100%) while the temperature was raised to -25°C. After three exchanges with 100% Lowicryl resin, each lasting 10 hours, polymerization was carried out using UV light at -25°C for 48 hours. This was followed by raising the temperature to 20°C and a further 48h of UV exposure.

A 3D confocal map of the resin block was then generated using the Zeiss LSM 780 NLO microscope. The block was mounted upside down on a drop of water in a glass-bottomed round dish and a 4X4 tiled scan was acquired using the 25X/0.8 NA multi-immersion objective. The cell of interest was identified and branded with the 2-photon laser set to a wavelength of 800 nm and 10% laser power. The block was then mounted onto a SEM stub using super-glue. A coating of silver paint was applied around the sa mple area, and the stub was gold-sputtered at 30 mA for 180s (Q150RS, Quorum).

### FIB-SEM acquisition

The whole-cell volume was acquired at an isotropic 10 nm voxel size at the Zeiss Crossbeam 540. Milling was done at 30 kV, 3 nA and SEM imaging at 1.5 kV accelerating voltage and current of 700 pA with an ESB detector (ESB Grid 1110V). The final dwell time for acquisition was 11 µs. The raw dataset was aligned using the Fiji plugin - Linear Stack Alignment with SIFT.

### Expansion microscopy sample preparation and imaging

hTERT-immortalized retinal pigment epithelial cells (RPE-1) were cultured according to standard condition, seeded on a 12 mm round coverslip, and fixed for 5min in -20 °C cold Methanol. Successively, samples were subjected to Ultra-Expansion microscopy (U-ExM) as described in ^24^. In brief, fixed samples were anchored by incubation in 2% Acrylamide / 1.4% Formaldehyde in PBS for 3h 30min at 37 °C, and incubated in a monomer solution containing 23% (w/v) sodium acrylate, 10% (w/v) acrylamide, 0.1% (w/v) N,N’-methylenebisacrylamide, 0.5% tetramethylethylendiamine and 0.5% ammonium persulfate in PBS for 1 h at 37 °C in a dark humidified chamber. For the homogenization, the polymerized gels were incubated for 15-30 min at room temperature (RT) on a rocker in denaturation buffer (50 mM Tris base, 200 mM SDS, 200 mM NaCl in water, pH 9) and then for 1 h 30 min at 95 °C, still in denaturation buffer in a thermomixer, before being expanded overnight in ddH2O. For immunostaining, expanded gels were incubated in PBS for 1 h at RT, blocked in a solution containing BSA 3%, Triton-X 100 0.05% in PBS for 30 min at 37 °C and stained overnight at 4 °C under constant agitation with the following primary antibodies: α-Tubulin (1:1500, Geneva antibody facility, ABCD_AA344), β-Tubulin (1:1500, Geneva antibody facility, ABCD_AA345) and acetyl-Tubulin-α (Lys40) (1:500, Thermo Fisher Scientific, 32-2700). Gels were then washed three times in PBS– 0.1% Tween for 30 min and stained for 2 h 30min at 37 °C under constant agitation with the following secondary antibodies: donkey anti-guinea pig Alexa Fluor 488 (1:1000, Jackson ImmunoResearch,706-545-148) and donkey anti-mouse Alexa Fluor 594 (1:1000, Thermo, A-21203). The nuclei were counter stained with Hoechst for 10 min at RT. Finally, stained gels were washed three times in PBS-0.1% Tween and expanded for 1 h at RT in ddH2O. The final expansion factor was measured to be 4.06. For imaging, small pieces of gels were mounted on 25 mm round precision coverslips (#1.5H) coated with poly-D-lysine and imaged on a Microscope Zeiss LSM 980 AIRY Fast (40 X objective,1.2NA).

## Acknowledgements

We thank Brady Johnston for help with Blender and the Advanced Light Microscopy Facility (ALMF) , the Advanced Mobile Laboratory (AML) and the Electron Microscopy Core Facility (EMCF) at the European Molecular Biology Laboratory (EMBL) in Heidelberg, Evident/Olympus, and Zeiss for their support; IT and HPC resources at the EMBL in Heidelberg for providing the essential computational infrastructure. We thank Jan Ellenberg for his feedback on our manuscript. This work was supported by the European Molecular Biology Laboratory, and by the Deutsche Forschungsgemeinschaft (DFG, German Research Foundation) - project number 452616889.

