## Supplementary Information. Supplementary note 1, Supplementary figure 1, and captions of supplementary videos. for "Microscopy Nodes: versatile 3D microscopy visualization with Blender"

Supplemental Video SV1. **Time-lapse render of a mitotic cell.** *Microscopy Nodes renders 5D microscopy stacks while combining render modes.* A mitotic cell with cell (yellow) and chromosome (purple) segmentations is shown, with a volumetric render of the fluorescent signal showing DNA (cyan). The grid is 10 µm.

Supplemental Video SV2. **Video of RPE1 cell expansion microscopy.** *Microscopy Nodes allows for animation of camera and visualization parameters.* The camera zooms in on the centrioles (acetylated tubulin, cyan) while the microtubules (αβ-tubulin, white) fade out. The nucleus, stained with Hoechst, is shown in pink. The acetylated tubulin subsequently fades out to replace with a purple model of the acetylated centriole, which is then appended with other centriolar protein elements (like PCM, subdistal appendages, and linker). The biological grid is 5 µm, the expanded grid is 20 µm.

Supplemental Video SV3. **Video showing a FIB-SEM dinoflagellate and its segmentations.** *Microscopy Nodes shows the context of volumetric EM by showing electron-sparse regions as transparent.* The render goes through a Z-stack showing separately sliced FIB-SEM volume and trichocyst segmentations (multicolor), to then rotate and show theca segmentation (blue), chloroplast segmentation (green), mitochondria segmentation (yellow).

Supplemental Note 1. **Volumetric render settings.** By default, Microscopy Nodes loads data with render settings to allow for quick and easy interaction. For some datasets, and especially in the light-scattering raytraced volume rendering often used for EM data, it may be useful to change some of these render settings. Note that changing these settings may require more computational power. Most of these settings are found in the "Render Properties" of Blender. The parameters include hyperparameters that define how many steps light rays maximally take to traverse a volume before failing (Render Properties > Volumes > Max Steps) and how many times a ray of light bounces (Render Properties > Light Paths > Transparent/Total/Volume). Additionally, for light-scattering volumes, one can also edit whether the light mostly backscatters or forward scatters (Volume Scatter > anisotropy, in the shader of the channel). Setting the anisotropy to negative values results in backscattering that highlights surface features, setting it to positive values produces forward scattering that penetrates deeper into the volume and can emphasize thin structures.

Supplemental Figure S1. **EM data filtered to membranes.** Dense 2D render using the same filter as shown in Fig 3C. Mapping to color in 2D rather than mapping to density in 3D makes this filter too strict in 2D, while it highlights the nuclear tunnels in 3D.

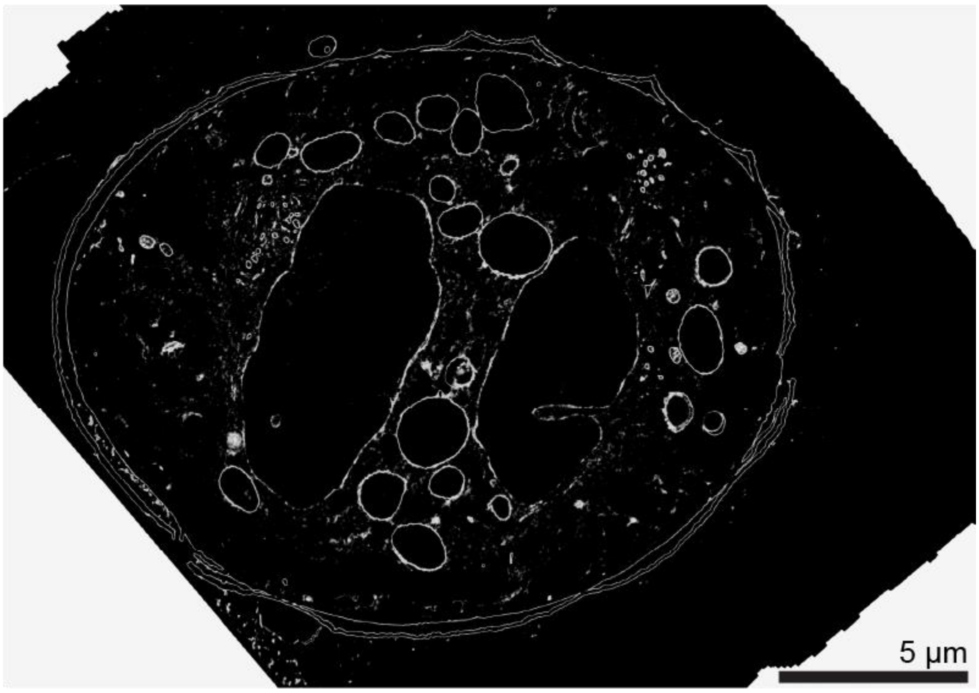
